# INDEX-db: The Indian Exome Reference database (Phase-I)

**DOI:** 10.1101/312090

**Authors:** Ahmed P Husayn, V Vidhya, Ravi P More, Mahendra S Rao, Biju Viswanath, Sanjeev Jain, Odity Mukherjee, ADBS Consortium

**Affiliations:** National Centre for Biological Sciences, Bengaluru, India; Institute of Bioinformatics and Applied Biotechnology, Bengaluru, India; Institute for Stem Cell Biology and Regenerative Medicine, Bengaluru, India; National Institute of Mental Health and Neuro Sciences, Bengaluru, India

**Keywords:** Population-specific database, Genetic variations catalogue, Indian population, Whole exome sequencing

## Abstract

Deep sequencing based genetic mapping has greatly enhanced the ability to catalog variants with plausible disease association. The bigger challenge now is to ascertain pathological significance to the array of identified variants to specific disease conditions. Differential selection pressure may impact frequency of genetic variations, and thus the detection of association with disease conditions, across populations. To understand the genotype to phenotype correlations, it thus becomes important to first understand the genetic variation spectrum of a population by creating a reference map. In this study, we report the development of phase I of a new database of coding variations, from the Indian population, with an aim to establish a centralized database of integrated information. This could be useful for researchers involved in studying disease mechanism at the clinical, genetic and cellular level.

Database URL: http://indexdb.ncbs.res.in

## Introduction

Human population has increased significantly in numbers across all geographical regions in the recent past, resulting in population specific genetic architecture. Such rapid population growth has significant impact on the occurrence and frequency of genetic variations, especially rare variants which may lie on conserved protein encoding sites, that may have a likely role in disease biology (Keinan and Clark, 2012). Next Generation Sequencing (NGS) strategies have greatly improved the ability to identify genetic variants, of varying frequencies. Recent studies to identify genetic variants associated with ‘common non-communicable disease’ suggest that these syndromes have high heritability, and that the risk arises from a polygenic contribution, caused by a combination of rare deleterious and common polymorphic modifier variants. NGS based evaluation of disease association thus becomes a useful way to identify disease genetic signature. A critical component of this analysis is the assignment of pathogenic relevance to the identified variants, done primarily by defining the frequency in affected individuals as compared to control, healthy samples. In this context, several genetic variation databases have been established incorporating different strategies and technological improvements (eg. haplotype mapping – HapMap project (The International HapMap Consortium, 2005); whole genome sequencing – 1000 Genomes project (The 1000 Genomes Project Consortium, 2015); whole exome sequencing – Exome Aggregation Consortium (Lek et al., 2016)). While the information gleaned from these databases improved our understanding of the complexities of the genetic architecture, it also reported that a significant proportion of genetic variations identified are population specific. We thus need a detailed evaluation in diverse populations, to better understand the genetic basis of epidemiology and semiology of human diseases by identifying modifier genetic variations (Bamshad et al., 2011; Craddock and Owen, 2010; Higasa et al., 2016; Hindorff et al., 2011).

The Indian subcontinent is estimated to see an increase in the number of individuals needing care for adult onset common disorders due to improved health care and life expectancy. Identification of disease specific genetic signature is a critical first step in identifying – a) disease associated genetic variations, b) molecular sub-typing of complex human phenotypes and c) at risk individuals with improved efficiency. A comprehensive reference variation map, established from a clinically normal cohort that is representative of this population, will be of great benefit. There have been several reports of cataloging genetic variation from the Indian population which have suggested presence of distinct genome level sub-structuring, and its probable impact on disease Biology (Narang et al., 2010; Rustagi et al., 2017; The HUGO Pan-Asian SNP Consortium, 2011; The Indian Genome Variation Consortium, 2005; Upadhyay et al., 2016). However, there are a few limitations to these studies – a) these predominantly catalogue germline variants; b) are designed to capture high frequency common variations, which is sufficient for deciphering population structure, but lack information on rare mutations, CNVs and disallow haplotype analysis; and, importantly c) are not available as open access reference map.

In this study we report the development and completion of phase 1 of a new accessible database-the INDian EXome database (INDEX-db), that catalogues variations in exonic and regulatory regions from healthy control individuals, across different geographical regions of southern India. To make the database a comprehensive resource for disease genetics studies, we have integrated WES derived SNV, CNV and phased LD information, along with expression data, on samples derived from a subset of individuals sequenced. We believe that such an integrated reference database may be valuable to understand the genomic architecture underlying susceptibility to disease, detect familial or geographical clustering of the population, and thus aid efforts to understand disease genetics.

## Materials and methods

### Samples information and ethical approval

Thirty one individuals tested to be asymptomatic for any adult onset common clinical illness were selected for the study at National Institute of Mental Health and Neuro Sciences, Bengaluru (Gender, age and other information of the individuals in Supplementary Table 1). The study was approved by the institutional ethics committee. Written informed consent was obtained from all participants prior to sampling. 10 ml of peripheral blood was collected under aseptic conditions and high molecular weight DNA isolated.

### Library preparation and exome sequencing

The genomic DNA was extracted from the blood and the Illumina Nextera Rapid Capture Expanded Exome kit was used for library preparation. Sequencing was carried out on Illumina Hiseq NGS platform. Quality check of the raw reads was performed using FASTQC tool (www.bioinformatics.babraham.ac.uk/projects/fastqc/). Only the paired-end raw reads with a score more than Q20 was filtered using Prinseq lite version 0.20.4 (Schmieder and Edwards, 2011) for further alignment to the reference genome. Reads were also checked for per base and per sequence quality scores, GC content, and sequence length distribution.

### Alignment and mapping of reads

The raw reads were aligned to the Human reference genome hg19 (GrCh37) using BWA tool version 0.5.9 (Li and Durbin, 2009). PCR duplicates in the mapped reads were marked using Picard (http://broadinstitute.github.io/picard/). INDEL realignment was performed using GATK version 3.6 (Depristo et al., 2011). Conversion of the sequence alignment file (SAM to BAM), indexing and sorting were done by samtools version 1.5 (Li et al., 2009). The quality check for the alignment on the mapped reads was performed using Qualimap version 2.2.1 (Okonechnikov et al., 2015).

### Detecting SNPs, indels and CNVs

SNPs and indels were called from the aligned files using Varscan2 version 2.3.9 (Koboldt et al., 2009; Koboldt et al., 2012) (with the criteria min coverage = 8, MAF >/= 0.25% and P </= 0.001). Depth of coverage was calculated using GATK version 3.8.0 (16) and this was used to detect copy number variations using XHMM (Fromer et al., 2012; Fromer and Purcell, 2014). XHMM employs principal component analysis to remove batch and target effects. Principal component analysis was performed on the entire read-depth matrix (31 individuals by 336,037 targets) and a hidden Markov model was applied to the normalized data to detect CNVs.

### Haplotype phasing

Haplotype pre-phasing was done for SNP genotypes from 31 individuals using SHAPEIT2 (v2.r837.GLIBCv2.12) (Delaneau et al., 2014; O’Connell et al., 2014). As a haplotype reference, we downloaded 1000Genome project Phase3 reference (http://mathgen.stats.ox.ac.uk/impute/1000GP_Phase3/) and used only the SAS subgroup haplotype reference. The phased data was visualized and haplotype blocks were generated based on the Dprime values computed for every comparisons between markers (SNPs) which are present within a distance range of 500kb using Haploview version 4.2 (Barrett et al., 2005). Default parameters were used which includes markers having MAF>0.05, p-value cutoff of 0.001, with maximum Mendelian errors of 1, minimum genotype percentage of 75%, exclusion of individuals with >50% of missing genotypes, with 95% confidence bounds (Gabriel et al., 2002).

### Development of INDEX-db

The SNPs and indels obtained from all the 31 individuals were merged using vcftools (Danecek et al., 2011) to create a merged SNPs and indels catalogue. This was annotated with ANNOVAR (reference assembly 65) (Wang et al., 2010). Copy number variations were pooled from all the individuals and used to create a reference copy number profile for the population. Pooling of data, functional analysis and other downstream analysis were performed using in-house shell and python scripts. The entire workflow of developing INDEX-db is shown in Fig. 1. The graphical genome browser for the database was developed on JBrowse version 1.12.3 (Skinner et al., 2009).

**Fig. 1:**
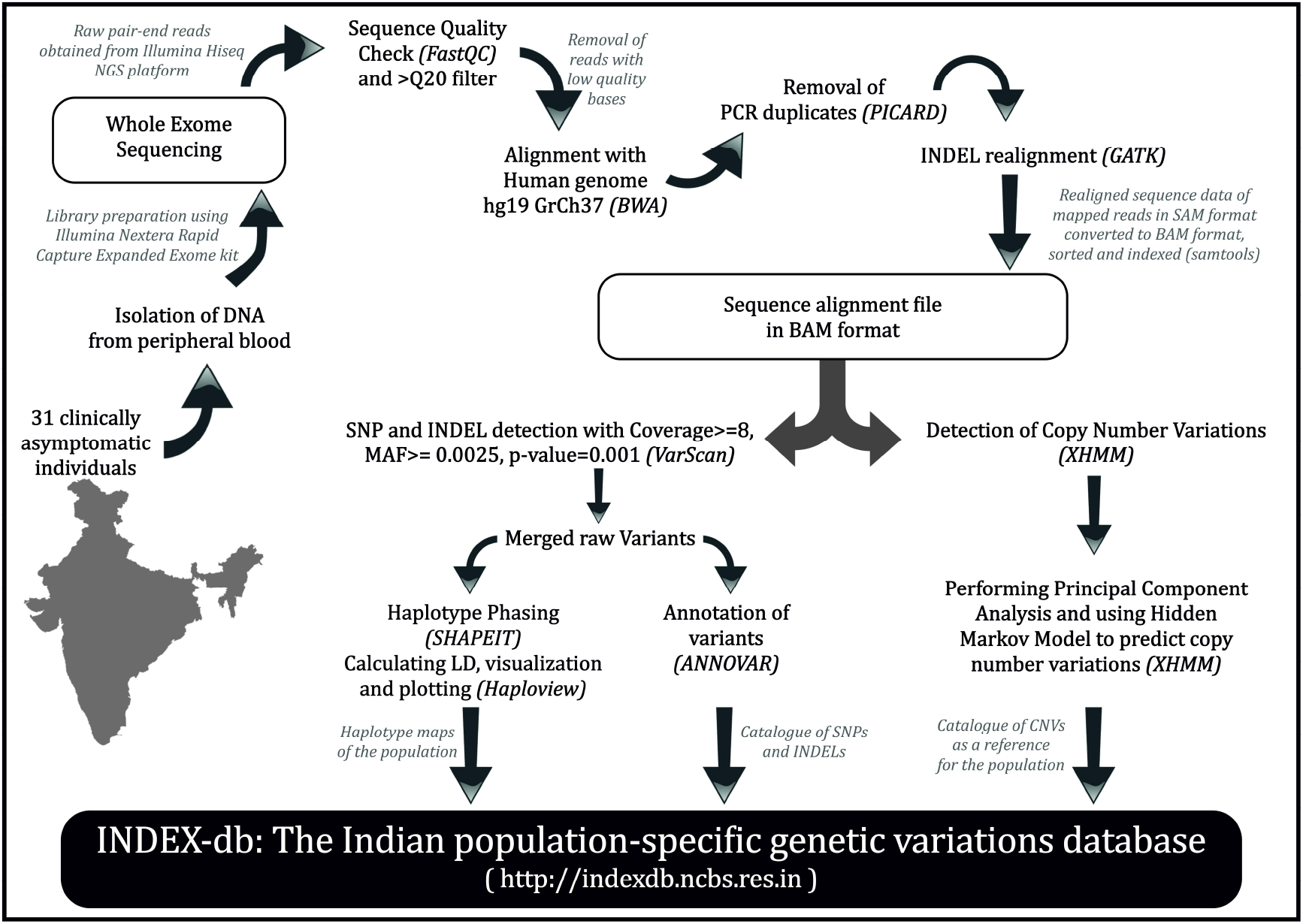
The workflow of the development of phase 1 of INDEX-db. The steps involved in the development of INDEX-db. The tools used in every step are mentioned in the brackets.

### Data availability

The raw sequence data has been deposited at the NCBI SRA database (SRA accession SRP135959). The entire database is hosted online at http://indexdb.ncbs.res.in and is freely accessible along with associated tools for querying and comparing user data to INDEX-db. The data is also available for download in standard formats at http://indexdb.ncbs.res.in/downloads.html. The SNPs are also deposited at the NCBI’s dbSNP (https://www.ncbi.nlm.nih.gov/SNP/snp_viewTable.cgi?handle=OMUKHERJEE_ADBS).

## Results

### INDEX-db: Variant summary profile

A total of 397,336 single nucleotide variations were identified in this phase 1 of the INDEX-db with an average 96% of the reads mapping to the reference genome at a mean coverage of 54.6% with at least 20X depths (Fig. 2). There was no significant bias seen, in terms of sequencing and/or sample QC (Fig. 2). About ~23% of the total genetic mutations identified were in the coding region, of which nearly half (51.34%) were a missense variation, followed by silent (43.36%), indel (1.8%), nonsense (0.81%) and splice sites (0.55%) (Fig. 3A). The ratio of non-synonymous (NS=49013) to synonymous variants (S=39876) was 1.23 (Fig. 2). The SNP profile observed in our study is comparable to exome sequencing reports published earlier (Lek et al., 2016; Rustagi et al., 2017; Upadhyay et al., 2016).

**Fig. 2:**
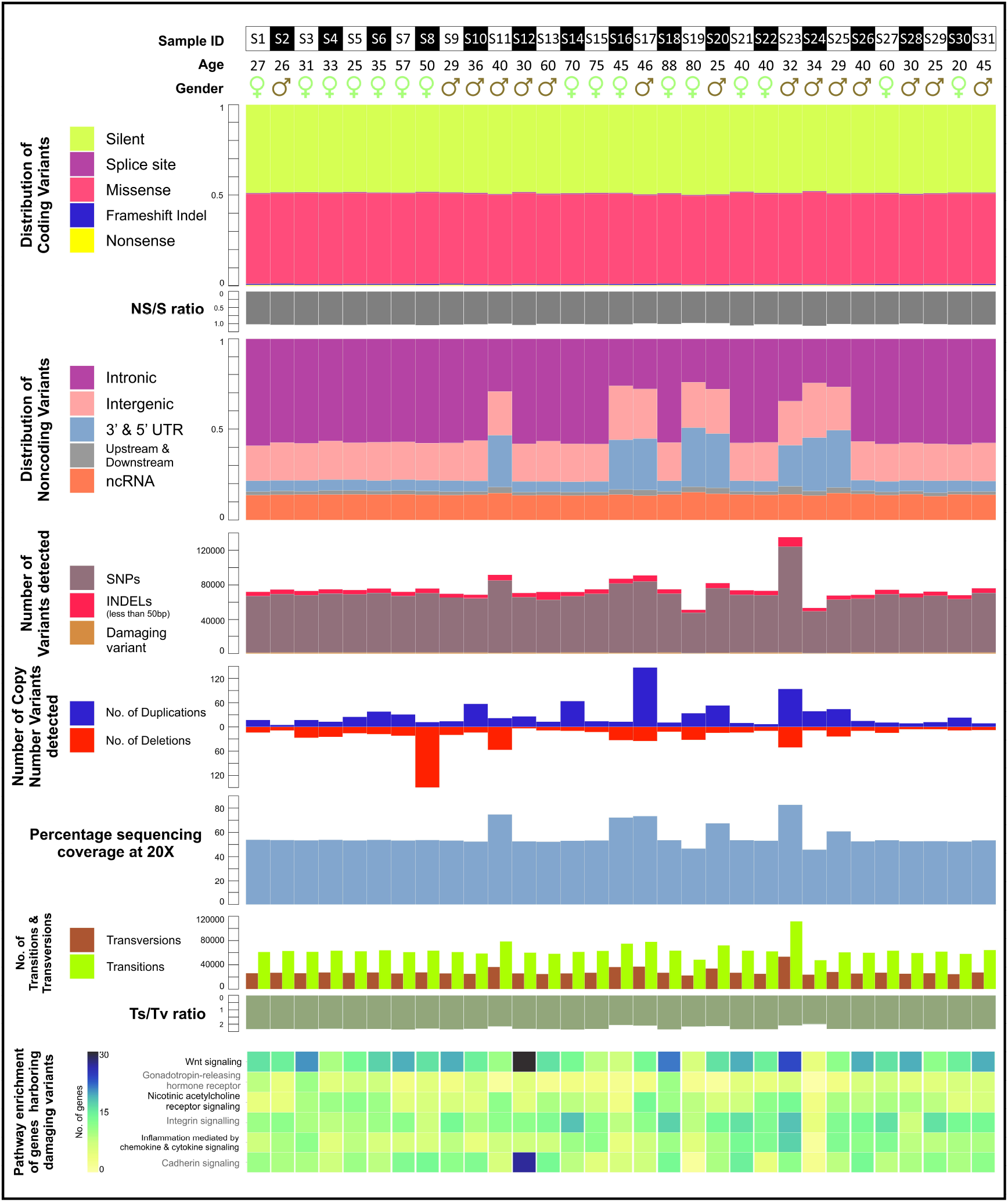
Variant summary profile. The distribution of coding and non-coding variants, the nonsynonymous to synonymous and the transitions to transversions ratios, and the percentage coverage of sequencing at 20X of 31 individuals catalogued in INDEX-db. The number of SNPs and CNVs detected in every individual with the pathway enrichment of genes harbouring damaging SNPs predicted by SIFT and PolyPhen2. Abbreviations: NS-Non-synonymous; S-Synonymous; UTR-Untranslated Region; ncRNA-Non-coding RNA; Ts-Transitions; Tv-Transversions.

**Fig. 3:**
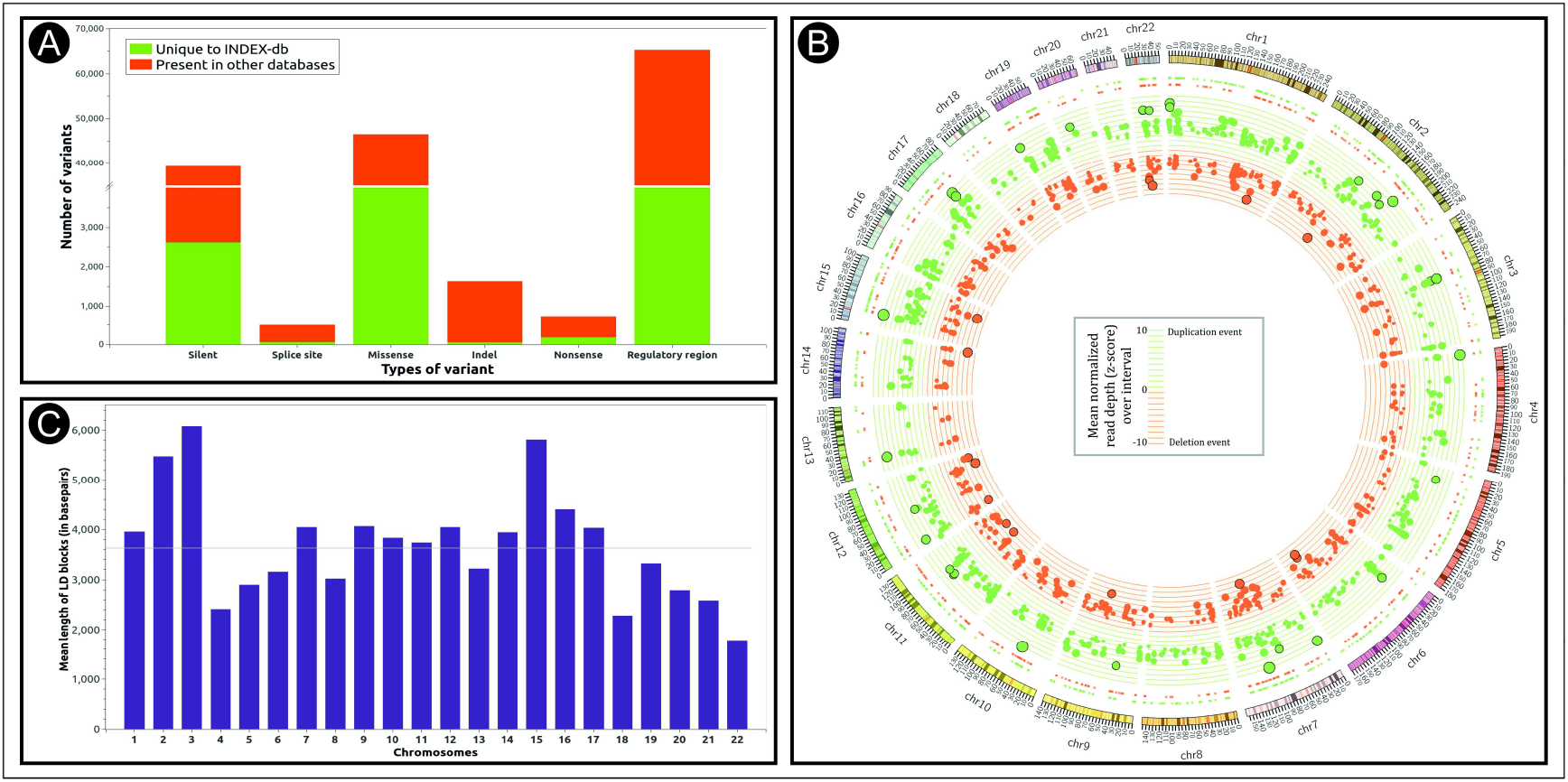
INDEX-db genetic catalogue. (A) Comparison of INDEX-db with other public databases. (B) The circos plot showing the copy number variation events catalogued in INDEX-db. The duplication and deletion events has been coloured green and red respectively. (C) Mean length of linkage disequilibrium blocks identified in autosomes.

Copy number variations (CNVs) contribute about one tenth of a percent of the total genetic variations of an individual, and it affects longer regions than both SNPs and/or short indels (The 1000 Genomes Project Consortium, 2015). The CNVs have a spectrum of phenotypic effects, from adaptive traits (Beckmann et al., 2007) to embryonic lethality (Hurles et al., 2008), and are implicated in many disorders including schizophrenia (Cook and Scherer, 2008), Down’s syndrome (Korenberg et al., 1994), kidney diseases (Nagano et al., 2018), diabetes (Ascencio-Montiel et al., 2017; Prabhanjan et al., 2016), hypertension (Boon-Peng et al., 2016; Marques et al., 2014), cancer (Araujo et al., 2014; Liu et al., 2013), bipolar disorder (Grozeva et al., 2013) etc. While array comparative genomic hybridization (aCGH) is considered as the standard for molecular assessment of genome-wide copy number detection (Pinkel et al., 1998; Pinkel and Albertson, 2005), methods have now been developed to detect copy number variations from NGS based exome and genome sequencing data (Yoon et al., 2009). Using a hidden Markov model, we identified a total of 1,538 CNVs in the size range of 50 bp to 3 mb in the INDEX-db phase I analysis represented as a circos plot (Fig. 3B). The number, size range and distribution of the detected CNVs in INDEX-db is comparable to other published data (MacDonald et al., 2014).

The common pattern in which variants are inherited across a population have critical importance in studying genetic correlates to rare and complex human diseases (The International HapMap Consortium, 2005). As parental genotype information may not be available for all the samples, reference phased haplotypes imputed using population relevant reference is valuable for disease genetics investigations. In INDEX-db phase I, we identified a total of 3365 LD blocks spread across the autosomes with an average block length of ~3.6 kb. (Fig. 3C).

To identify if there exists any population specific mutation/recombination hotspots, we computed the mean density of the variants (SNPs and CNVs) and the resulting LD blocks across the chromosomes by calculating the number of variants and/or LD blocks per million base pairs for each chromosome (normalizing for chromosome size). Apart from chromosome 19 which showed a significantly higher number of SNPs and CNVs, we did not observe any significant clustering of variants in any other chromosome (Fig. 4A-B). The increased density of variants observed in chromosome 19 may be due to the highest gene density or the presence of many paralogues of immunoglobulins localized to this chromosome that may undergo repeated duplication and/or mutation as reported by earlier studies (Castresana, 2002; Grimwood et al., 2004). We also observed that chromosome 19 had the highest number of LD blocks compared to the other chromosome. This could be due to the increased number and polymorphic diversity of the variants localized to this region resulting in low levels of common haplotypes. (Fig. 4C and Supplementary Table 2).

**Fig. 4:**
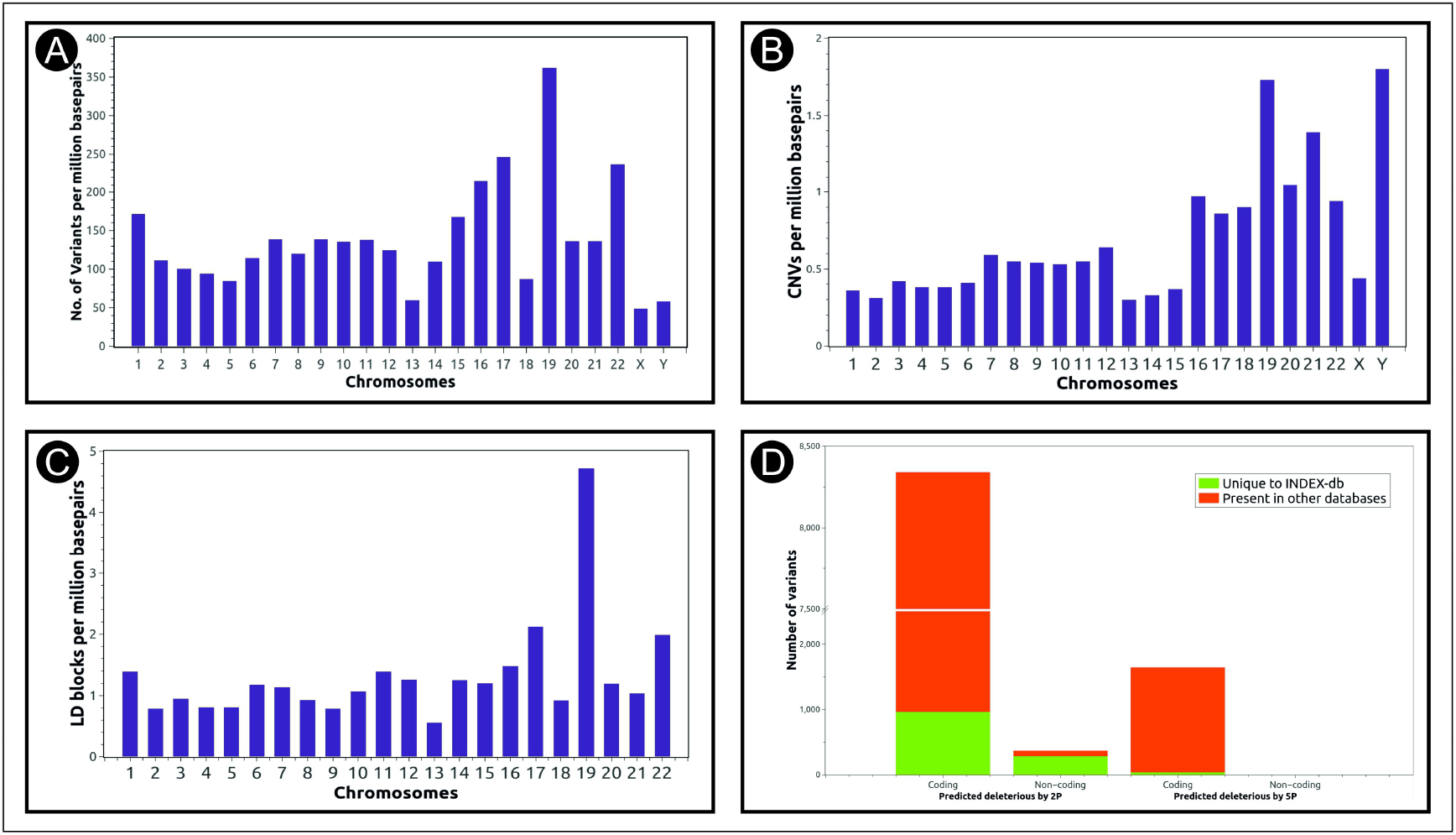
The distribution of genetic variations across chromosomes. The number of (A) SNPs, (B) CNVs and (C) LD blocks catalogued in INDEX-db per million basepairs in each chromosome. (D) In silico prediction of deleteriousness on coding and non-coding variants by 2 predicting algorithms (SIFT and PolyPhen2) and 5 algorithms (SIFT, PolyPhen2, Mutation Taster, Mutation Accessor, LRT predictor).

Population genetics studies have shown that there is greater genetic drift in East Asian populations, impacting the individual mutation burden load (Balick et al., 2015; Gao and Keinan, 2016; Simons and Sella, 2016). To ascertain the value of INDEX-db as a reference resource for disease genetics studies for the Indian population, we compared the INDEX-db phase I data with two publicly available databases. We used the ExAC as it is one of largest exome sequencing reference database with significant representation of South Asian population (although low representation from the pan-Indian population), and the AP-SAS, as it is a WGS based resource generated using samples from southern India. The variant parameter profile identified in INDEX-db are comparable to these other databases (Supplementary Table 3).

We found 12% (48732) of the variants identified were unique to INDEX-db phase I (Fig. 3A, Supplementary Table 3). Within the coding region, this translated to 8860 (~2.23%) variations, out of which 966 had a functional annotation of being ‘deleterious’ by two *in silico* algorithms (Fig. 4D) (Adzhubei et al., 2010; Ng and Henikoff, 2003). We found ~20% of coding variants identified in INDEX-db to be common between ExAC, and ~ 7% common to AP-SAS. The observation of low overlap between INDEX-db and AP-SAS could be attributed to the low coverage in the whole genome sequencing design of the AP-SAS study (~2X mean coverage). Differences between ExAC-SAS and INDEX-db could perhaps be attributed to population specific variation signature, especially since ExAC-SAS has a low representation from the Indian population. The mutational profile obtained in the phase I of this database is comparable to other databases, though currently limited by the number of individuals it represents.

### Functional relevance of INDEX-db as a population specific reference database for disease association studies

Based on the protein perturbing impact, mutations can be classified as benign (tolerated), deleterious (loss of function) and neutral (no influence). Functional impact of genetic variants identified in INDEX-d, was analyzed using SIFT (Ng and Henikoff, 2003) and PolyPhen2 (Adzhubei et al., 2010) algorithms that predict the functional consequences based on conservation and protein structure. A total of 8345 non-synonymous variants (9% of the total protein-coding variants) spanning 5097 genes were predicted to be damaging by SIFT and Polyphen2 in INDEX-db (Fig. 4D). Of these, 11% (966) spanning 700 genes were novel to INDEX-db. The percentage of damaging and unique variants observed in our dataset is in the range as reported earlier, although the specific genes and/or pathways may differ (The 1000 Genomes Project Consortium, 2015). To evaluate the biological impact of these deleterious mutations, we performed enrichment analysis on the genes harbouring deleterious mutations, and found increased enrichment for Wnt signaling, Nicotinic acetylcholine receptor signaling, Integrin signalling, cytokine signaling and Cadherin signalling pathway (Fig. 2). These pathways are implicated in various disorders including severe mental illness, diabetes and cancer. (Supplementary Table 4). Assessing the frequency of these rare variants in the general, healthy population may be useful to understand the genetic contributions to risk for disease, and also the relation with particular clinical syndromes. The comprehensive profile of genetic variations cataloged in INDEX-db phase I is detailed in Fig. 2.

## Discussion

We report the development of a new database, INDEX-db, which summarizes variations in coding and regulatory regions, identified from healthy control individuals. The first phase of the database consists of 31 individuals from southern India. We have integrated the exome sequencing results with expression data generated from a subgroup of the individuals constituting INDEX-db. The database is also layered with information regarding CNVs and phased LD mapping. The integrated database is available freely at http://indexdb.ncbs.res.in along with associated tools for querying and comparing user input data to INDEX-db.

The INDEX-db is in its first phase and thus in comparison to other public databases is limited in terms of the number of individuals sequenced to represent the population, but the variant profile we report in our pilot phase is comparable to population-based databases signifying its value in terms of giving population-specific information.

Genetic basis of complex disorders need to be better understood in India where the burden of these disorders is expected to increase significantly in the coming decades. In this context, we believe that an integrated reference database may be valuable to understand the genomic architecture underlying susceptibility to disease, familial or geographical clustering of the population and aid in disease genetics studies.

## Acknowledgements

The authors are grateful to all the volunteers who participated in the study. We thank Drs. Lakshmi Narayanan Kota, Manasa Seshadri and Ravi Kumar Nadella for recruitment of the control individuals, their clinical assessments and initial sample processing. Ten individuals were recruited as part of a Center of excellence grant in collaboration with Geriatric clinic team of NIMHANS (Profs Mathew Varghese, Sivakumar PT and other clinical staff). The authors would like to thank the sequencing core facility at IGIB (Dr. Faruq Mohammed) and NCBS (Dr. Awadhesh Pandit) for sample processing and data generation. The authors would like to thank all investigators of ADBS consortia for providing valuable inputs to the study and the manuscript.

The ADBS consortium members: Biju Viswanath^#^, Naren P. Rao^#^, Janardhanan C. Narayanaswamy^#^, Palanimuthu T Sivakumar^#^, Arun Kandaswamy^#^, Muralidharan Kesavan^#^, Urvakhsh Meherwan Mehta^#^, Ganesan Venkatasubramanian^#^, John P. John^#^, Odity Mukherjee^@^, Meera Purushottam^#^, Ramakrishnan Kannan^#^, Bhupesh Mehta^#^, Thennarasu Kandavel^#^, Binukumar B.^#^, Jitender Saini^#^, Deepak Jayarajan^#^, Shyamsundar A.^#^, Sydney Moirangthem^#^, Vijay Kumar G.^#^, Jagadisha Thirthalli^#^, Prabha S. Chandra^#^, Bangalore N. Gangadhar^#^, Pratima Murthy^#^, Mitradas M. Panicker^*^, Upinder S Bhalla^*^, Sumantra Chattarji^*@^, Vivek Benegal^#^, Mathew Varghese^#^, Janardhan YC Reddy^#^, Padinjat Raghu^*^, Mahendra Rao^@^, Sanjeev Jain^#^.

\# National Institute of Mental Health and Neuro Sciences (NIMHANS), Bengaluru, India

* National Centre for Biological Sciences – Tata Institute of Fundamental Research (NCBS – TIFR), Bengaluru, India

@ Institute for Stem Cell Biology and Regenerative Medicine (InStem), Bengaluru, India

## Conflicts of interest

There is no conflict of interest.

## Funding Source

The study was supported by government funded research grant under the aegis of Department of Biotechnology (grant number BT/PR17316/MED/31/326/2015) and Pratiksha trust. Ten individuals were recruited as part of a Center of excellence grant from Department of Biotechnology (grant number BT/01/CEIB/11/VI/1). HAP is supported by a grant from the Department of Biotechnology (grant number BT/PR12422/MED/31/287/2014). The funding agencies had no role in study design; in the collection, analysis and interpretation of data; in the writing of the report; and in the decision to submit the article for publication.

## Supplementary data

**Supplementary Table 1:** Sample IDs, age and gender of the individuals of INDEX-db

**Supplementary Table 2:** Mean density of the variants (SNPs and CNVs) and the resulting LD blocks across the chromosomes

**Supplementary Table 3:** Comparison of INDEX-db with other public databases

**Supplementary Table 4:** Deleterious variants in genes (known to be implicated in a spectrum of diseases) identified in the samples and comparison with public databases

